# Analysis of genomic-length HBV sequences to determine genotype and subgenotype reference sequences

**DOI:** 10.1101/831891

**Authors:** Anna L McNaughton, Peter Revill, Margaret Littlejohn, Philippa C Matthews, M Azim Ansari

## Abstract

Hepatitis B virus (HBV) is a diverse, partially double-stranded DNA virus, with 9 genotypes (A-I), and a putative 10^th^ genotype (J), thus far characterised. Given the broadening interest in HBV sequencing, there is an increasing requirement for a consistent, unified approach to HBV genotype and subgenotype classification. We set out to generate an updated resource of reference sequences using the diversity of all genomic-length HBV sequences available in public databases. We collated and aligned genomic-length HBV sequences from public databases and used maximum-likelihood phylogenetic analysis to identify genotype clusters. Within each genotype, we examined the phylogenetic support for currently defined subgenotypes, as well as identifying well-supported clades and deriving reference sequences for them. An alignment of these reference sequences and maximum-likelihood phylogenetic trees of the sequences are provided to simplify classification. Based on the phylogenies generated, we present a comprehensive set of HBV reference sequences at the genotype and subgenotype level.

## INTRODUCTION

Hepatitis B virus (HBV) is the prototype virus of the *hepadnaviridae* family, an unusual family of partially double-stranded (ds)DNA viruses approximately 3.2kb in length encoding four genes, including a reverse transcriptase, encoded in overlapping reading frames on a circular genome (1,2). To date, 9 genotypes of HBV have been characterised (A-I), along with a putative 10^th^ genotype (J) (3). The viruses display a relatively high amount of variation for a dsDNA virus (4), with the viral genotypes being further subdivided into upwards of 30 subgenotypes, many of which have distinct geographic and clinical associations (2).

At present, HBV genotyping is not routinely being performed in most clinical settings, as it has not been widely considered as relevant to patient management. However, as more treatment data become available and improvements are made in patient-stratified clinical care, guidelines may change to reflect different genotype-specific recommendations. This has been exemplified by the management of hepatitis C virus infection (5), and a similar approach for HBV is starting to emerge, with recent EASL guidelines for HBV treatment suggesting different stopping points for treatment non-response in genotypes A-D (6). Evidence increasingly supports a role for HBV genotype in influencing disease progression including risk of developing chronic infection, e-antigen seroconversion, transmission mode, and the development of hepatocellular carcinoma (7,8). Studies often refer to ‘wild-type’ virus (8,9), but ‘wild-type’ for one genotype may not reflect consensus for other genotypes (10,11). Understanding the diverse range of HBV strains circulating globally, and their associations with disease will allow us to move towards a more specific and precise approach to analysis.

A consistent approach to HBV subtyping is increasingly relevant, as interest in studying the associations between disease outcomes and viral sequence expands, and deep sequencing is used to investigate viral diversity in increasingly larger cohorts. A progressive body of sequencing data is emerging, and this expansion of available data is likely to increase over time (Suppl. Fig 1); however, the number of sequences covering the full genome lag a long way behind sequences for single genes or shorter fragments. To date, numerous subtyping misclassifications have been documented for HBV, predominantly driven by the use of partial genome sequences rather than full-length genomes (9,10), the inappropriate classification of recombinant strains (11), and publications re-designating same subgenotype classifications to distinctly different strains (12,13). Whilst a number of HBV sequence and analysis resources already exist, including HBVdb (14), HBVRegDB (15), HBVDR (16) and geno2pheno (https://hbv.geno2pheno.org/), each defines different sets of reference sequences, and frequently only at the genotype level. Despite proposed criteria for defining HBV genotypes and subgenotypes (3), currently there is no single resource where all the necessary sequences and their designation is readily available for use.

With this dataset, we have set out to provide a unified resource, taking a pragmatic approach to describing the phylogeny of HBV genotypes A-I using all available genotyped sequences from the HBVdb repository (14). We removed highly similar sequences from downloaded alignments and performed maximum-likelihood phylogenetic analysis. We used the phylogeny of whole genome sequences to identify well-supported clades and selected representative reference sequences for each identifiable subgenotype.

## RESULTS

### Generation of a dataset of genomic-length HBV sequences

We downloaded a total of 7108 full length HBV sequences from the hepatitis B virus database (HBVdb, https://hbvdb.ibcp.fr/HBVdb/) on 31st of January 2019. Following clean-up of the dataset to remove identical sequences and recombinants (as described in detail in the methods section) we were left with a total of 5972 sequences. We aligned the sequences using MAFFT and calculated pairwise distances for hierarchical clustering, which was then used to remove highly similar isolates (see methods). At this stage we were left with 2839 sequences on which we based our phylogenetic analysis (all sequences shown in Fig. 1).

**Figure 1.**
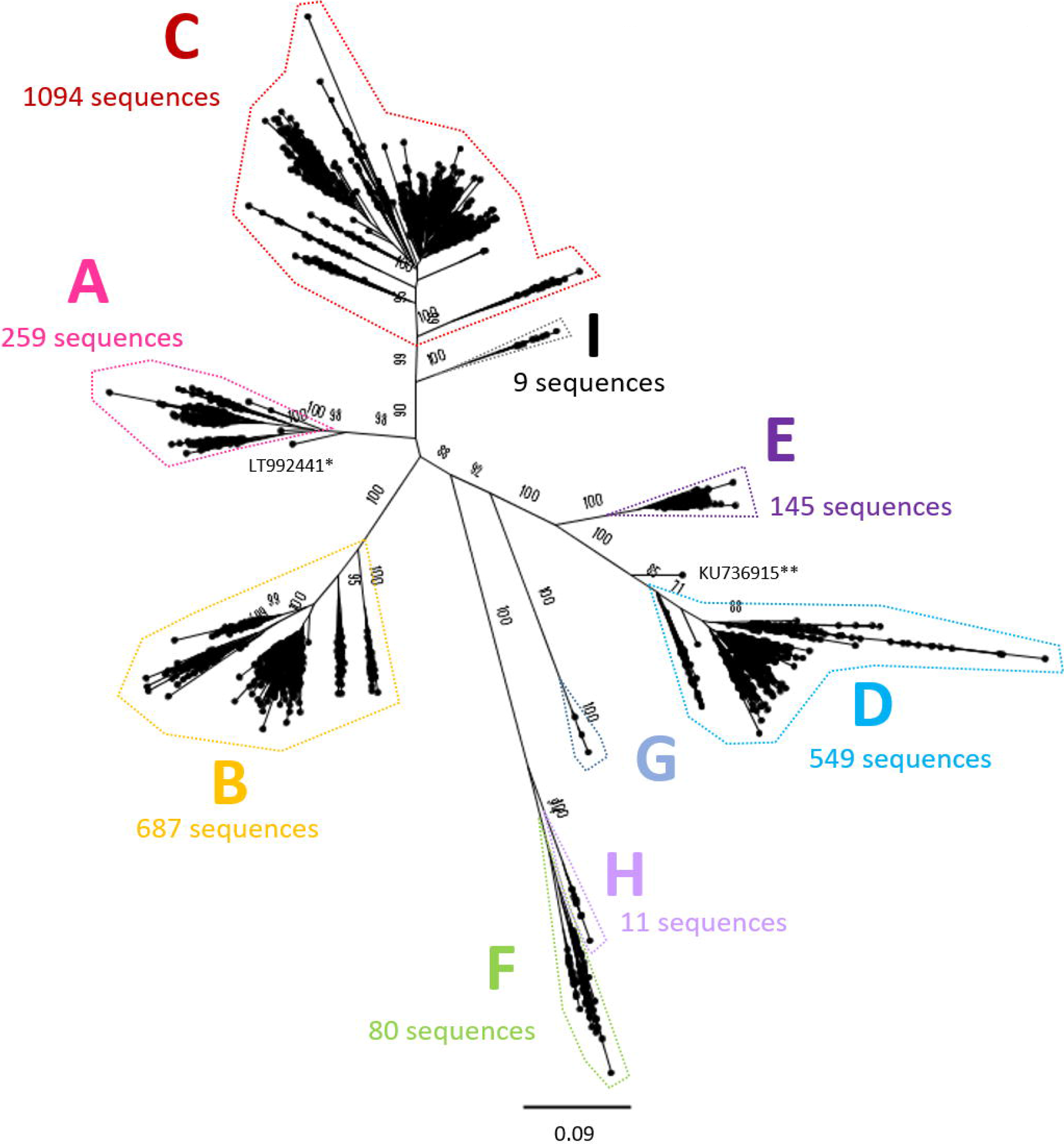
Genomic-length maximum-likelihood phylogeny of all full length genotype A-I HBV sequences included in analysis (n=2839), indicating the number of sequences in each genotype analysed separately in Figures 2–9. Maximum-likelihood phylogenetic analysis of sequences used in this study after removing highly similar sequences, demonstrating the strong genotypic clustering of the sequences. Bootstrap support ≥70 after 1000 replicates is given for the deepest branches on the tree. The scale bar indicates the estimated nucleotide substitutions per site. *A strain known to be from a 14^th^ century skeleton clustering distantly with genotype A, LT992441, was removed from the analysis. ** KU736915 was identified as a genotype D/E recombinant and removed from the subsequent analysis.

### Classification of HBV genotypes, subgenotypes and designation of references

We selected appropriate reference sequences to represent each genotype and subgenotype groups; these sequences are shown in the context of the overall phylogeny highlighted in Figures 2–9 and listed in Table 1. In cases where we could identify multiple distinct clades within a single subgenotype (subgenotypes A1, B2, C2, D1; Figs 2–5), we selected a reference for each clade to ensure the diversity within the subgenotype was represented. In genotypes A-D, the most well-represented subgenotypes contained ≥45% of all the sequences within that genotype. Sequences were most evenly distributed between subgenotypes in genotype F, with the commonest subgenotype accounting for 33% of sequences.

**Table 1.**
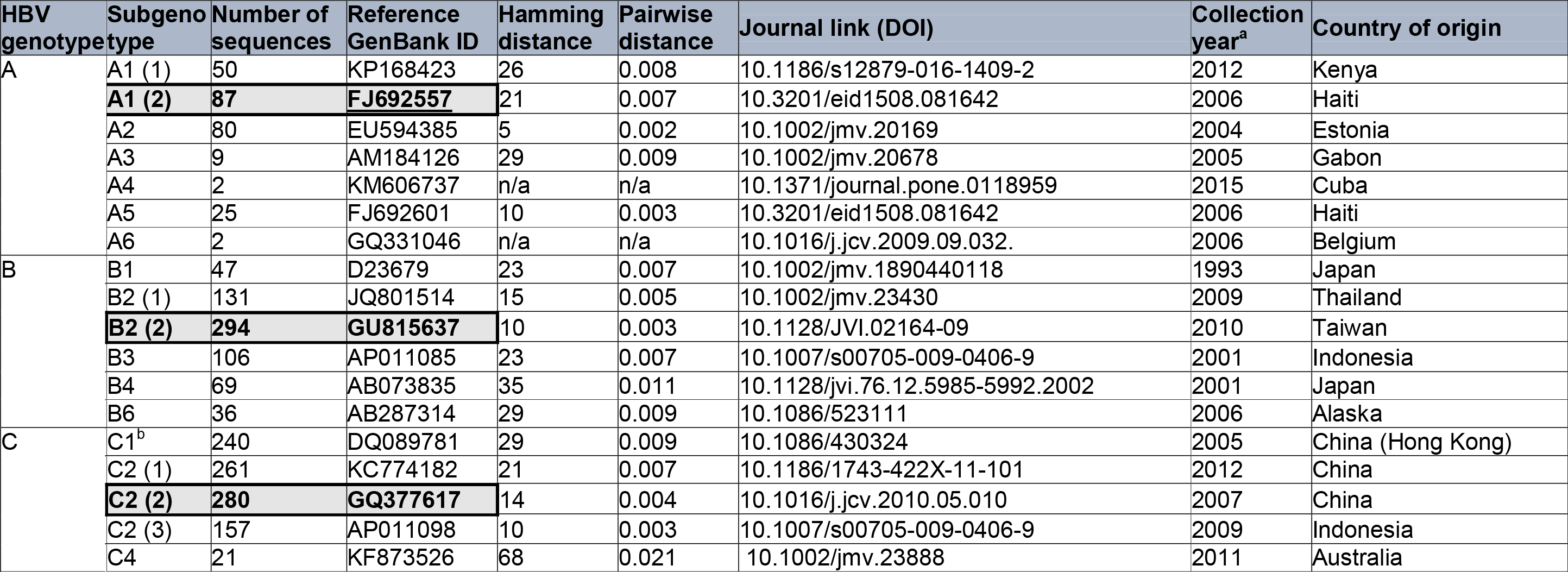
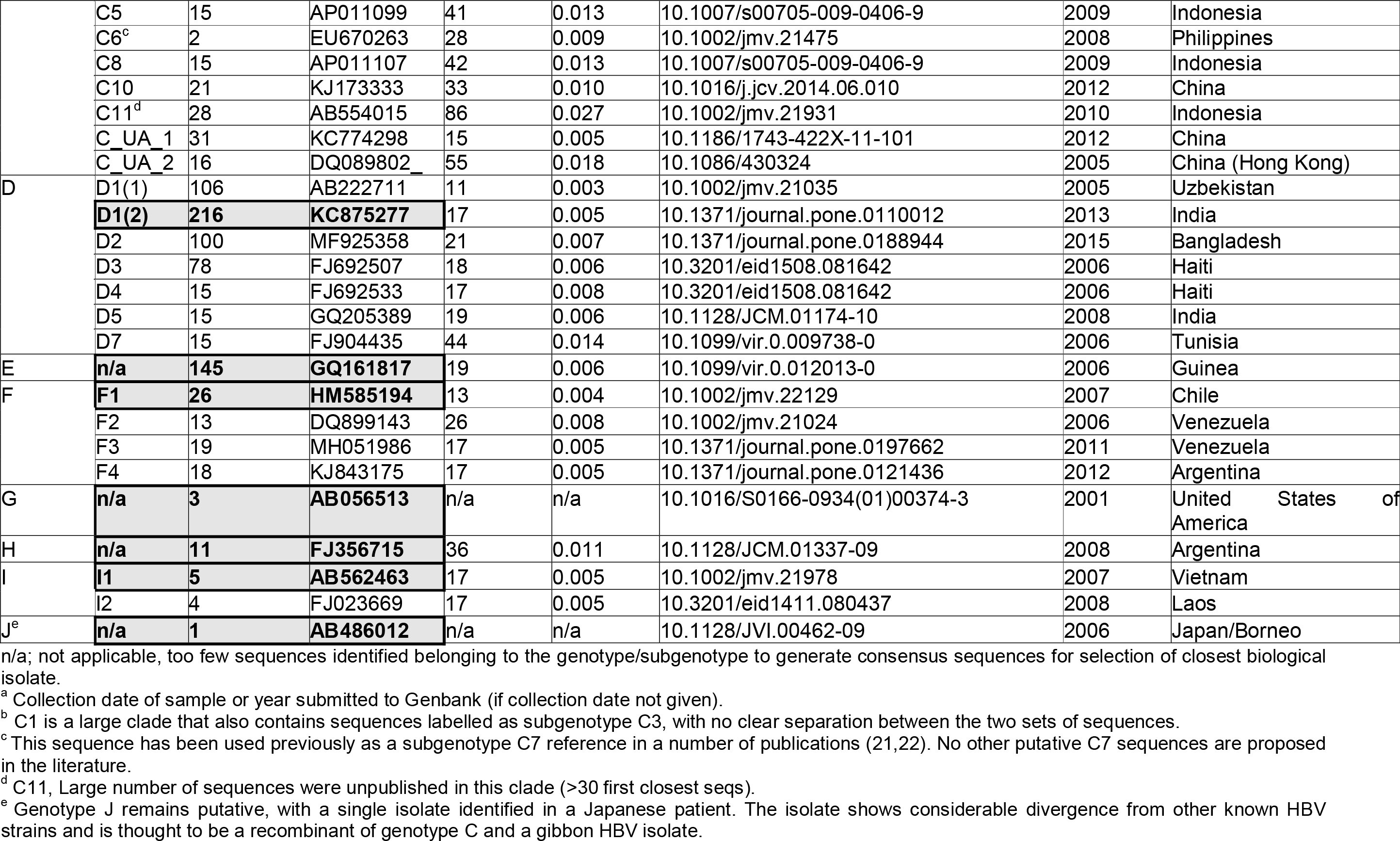
Proposed reference sequences for HBV genotypes, subgenotypes and clades. The number of sequences in each clade is given for each subgenotype and clade identified. Note that the total sum of the subgenotype sequence clusters may not correspond to the total number of genotype sequences, as a number of sequences did not group within a specific clade. Reference sequences for the genotypes are highlighted in grey boxes. In Figures 2–9, genotype references are marked with blue dots and subgenotype reference sequences are marked with red dots. Subgenotypes A4, A6, B5, C7, C9 and D6 are not included as these sequences either did not cluster as monophyletic clades or were not retained in our analysis. Hamming distance indicates the number of nucleotide differences between the clade consensus and the chosen reference. The pairwise distance is the number of nucleotide differences between the clade consensus and the chosen reference normalised by length of the genome.

**Figure 2.**
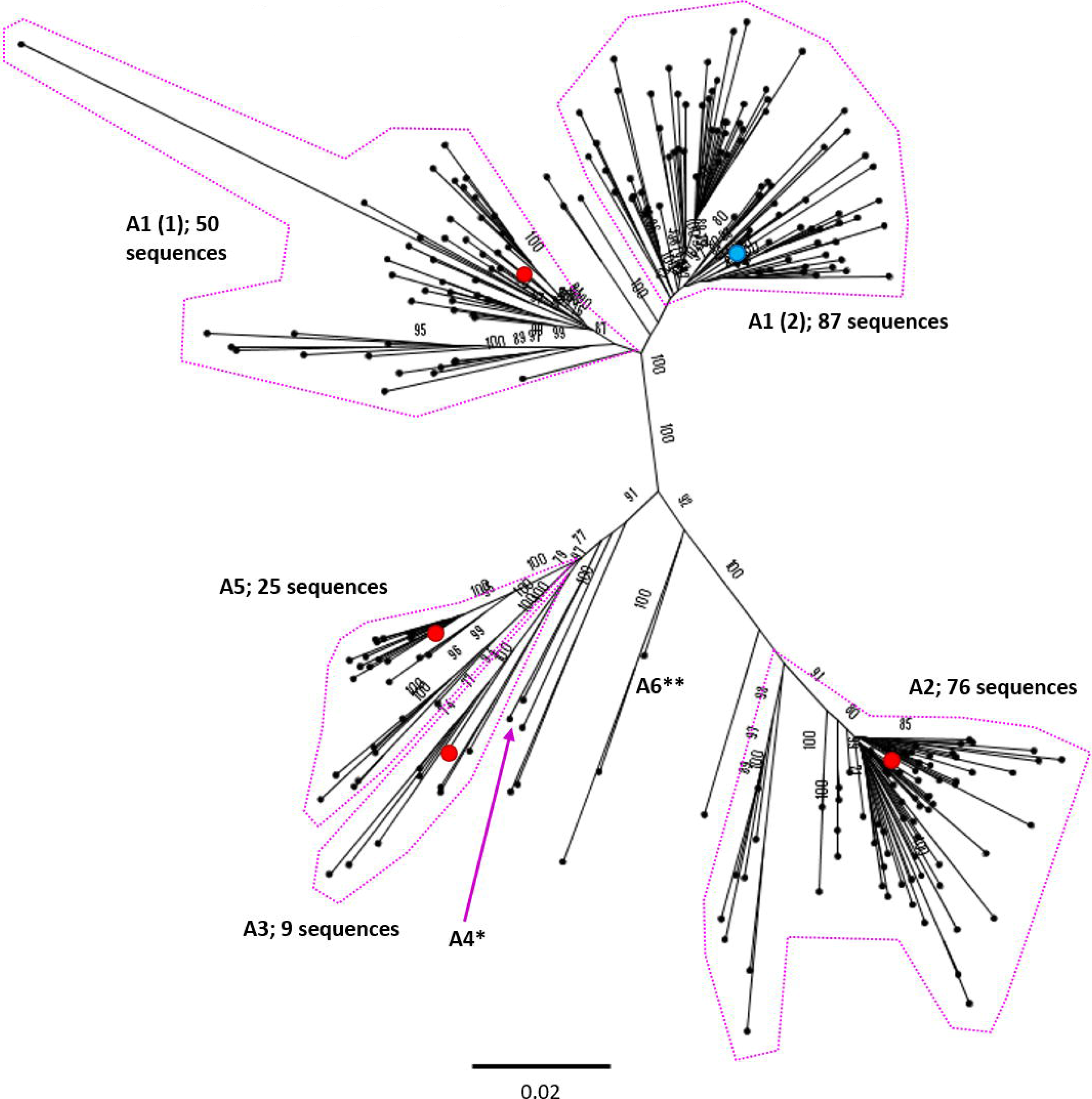
Genomic-length maximum-likelihood phylogeny of HBV genotype A sequences (n=259). Well-defined clades have been highlighted with coloured dotted lines and reference sequences for each clade indicated (red dots). Proposed reference strain for the genotype, FJ692557 is highlighted with a blue dot. The subgenotype is given where it could be reliably identified. Bootstrap support for branches ≥70 after 1000 replicates is indicated. The scale bar indicates the estimated nucleotide substitutions per site. Previous work has confirmed that there are at least 5 genotype A subgenotypes, although debate continues about whether or not A3, A4 and A5 should all be considered ‘quasi-subgenotype A3’ (9). Few sequences for subgenotypes *A4 (KM606737) and **A6 (GQ331046) were retained in the study after pairwise analysis. The putative subgenotype A6 has previously been identified in three African-Belgian patients (27).

**Figure 3.**
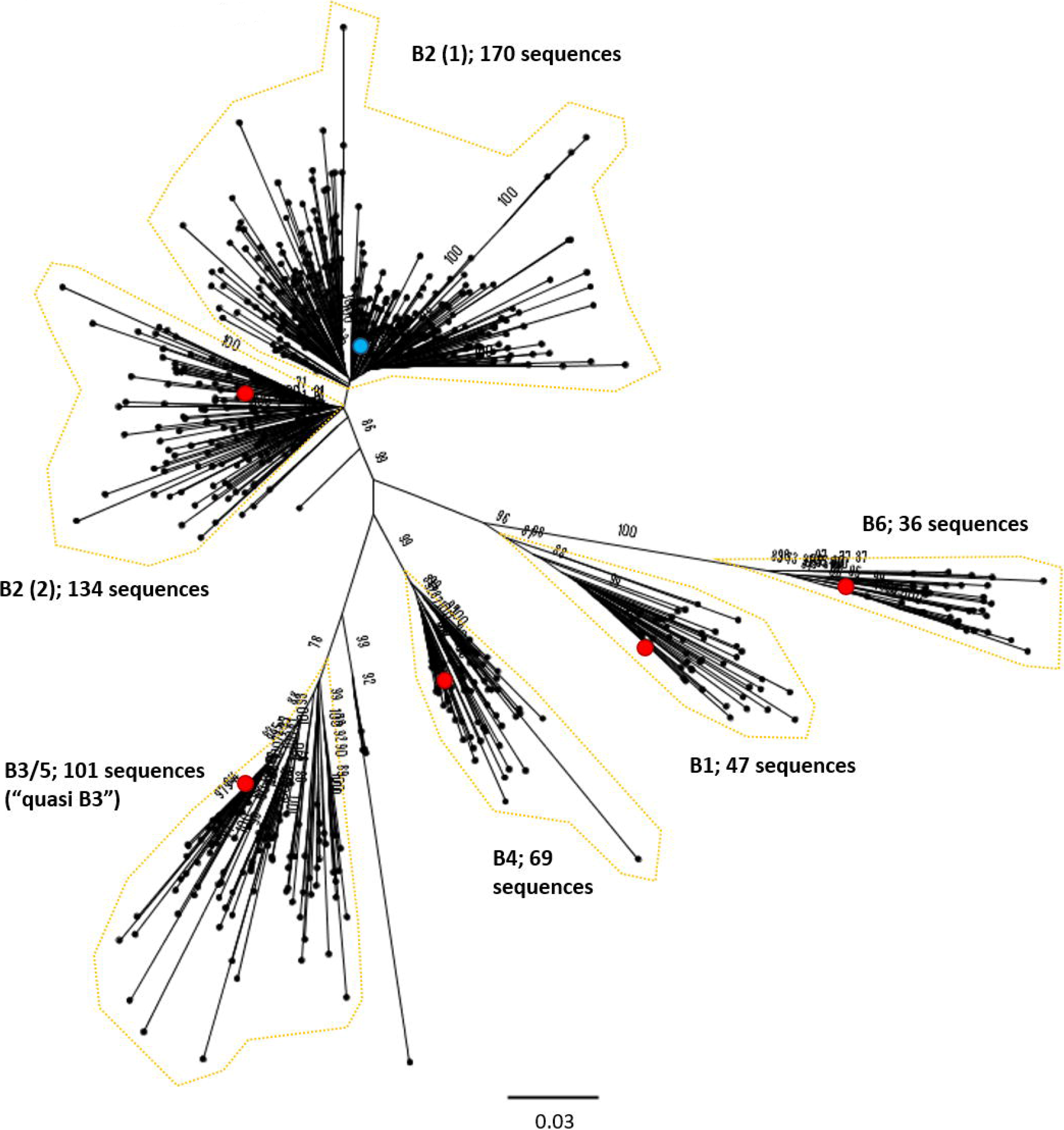
Genomic-length maximum-likelihood phylogeny of HBV genotype B sequences. Well-defined clades have been highlighted with coloured dotted lines and reference sequences for each clade indicated (red dots). Proposed reference strain for the genotype, GU815637 is highlighted with a blue dot. The subgenotype is given where it could be reliably identified. Bootstrap support for branches ≥70 after 1000 replicates is indicated. The scale bar indicates the estimated nucleotide substitutions per site. An evaluation of the genotype B phylogeny reclassified a number of putative subgenotypes as quasi-B3, with debate continuing on whether or not this should also include B5 (50).

**Figure 4.**
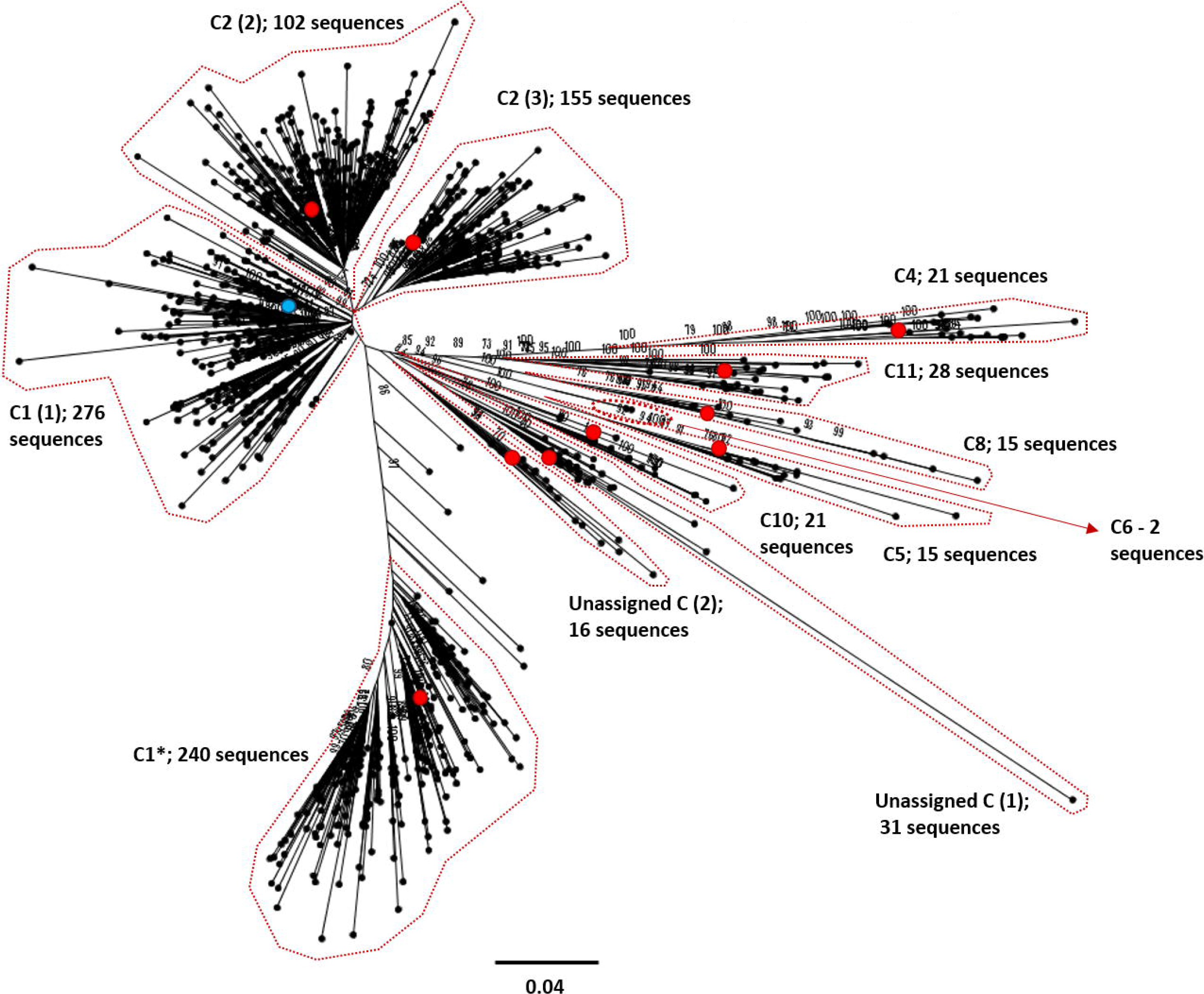
Genomic-length maximum-likelihood phylogeny of HBV genotype C sequences. Well-defined clades have been highlighted with coloured dotted lines and reference sequences for each clade indicated (red dots). Proposed reference strain for the genotype, GQ377617 is highlighted with a blue dot. The subgenotype is given where it could be reliably identified. Bootstrap support for branches ≥70 after 1000 replicates is indicated. The scale bar indicates the estimated nucleotide substitutions per site. We were unable to verify the subgenotype of two genotype C clades, and these have been designated unassigned clade 1 and 2 (unassigned_C (1) and unassigned_C (2), respectively).

**Figure 5.**
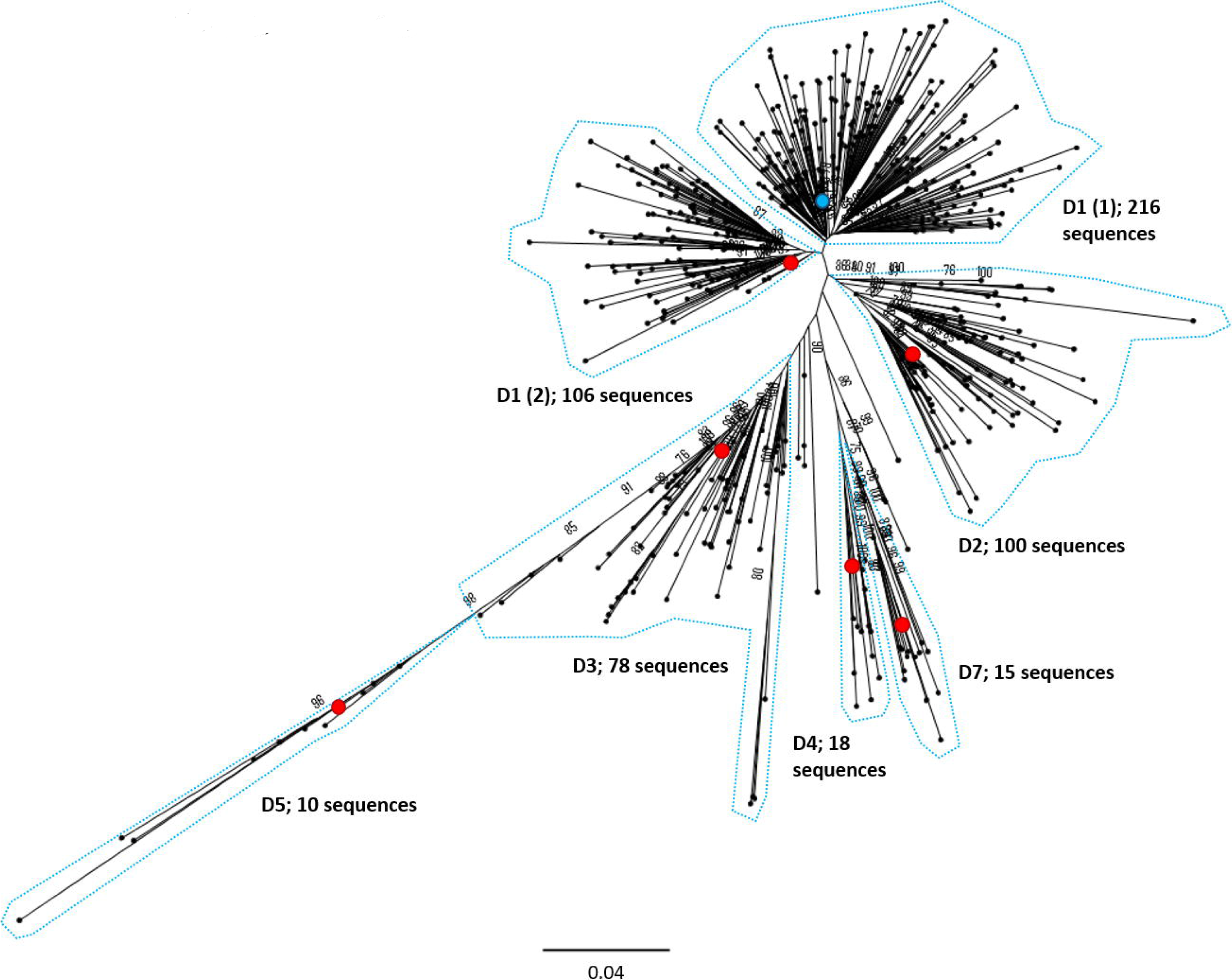
Genomic-length maximum-likelihood phylogeny of HBV genotype D sequences. Well-defined clades have been highlighted with coloured dotted lines and reference sequences for each clade indicated (red dots). Proposed reference strain for the genotype, KC875277is highlighted with a blue dot. The subgenotype is given where it could be reliably identified. Bootstrap support for branches ≥70 after 1000 replicates is indicated. The scale bar indicates the estimated nucleotide substitutions per site. Previous work has indicated D3 and D6 strains cluster together and should be classed as a single subgenotype (51).

**Figure 6.**
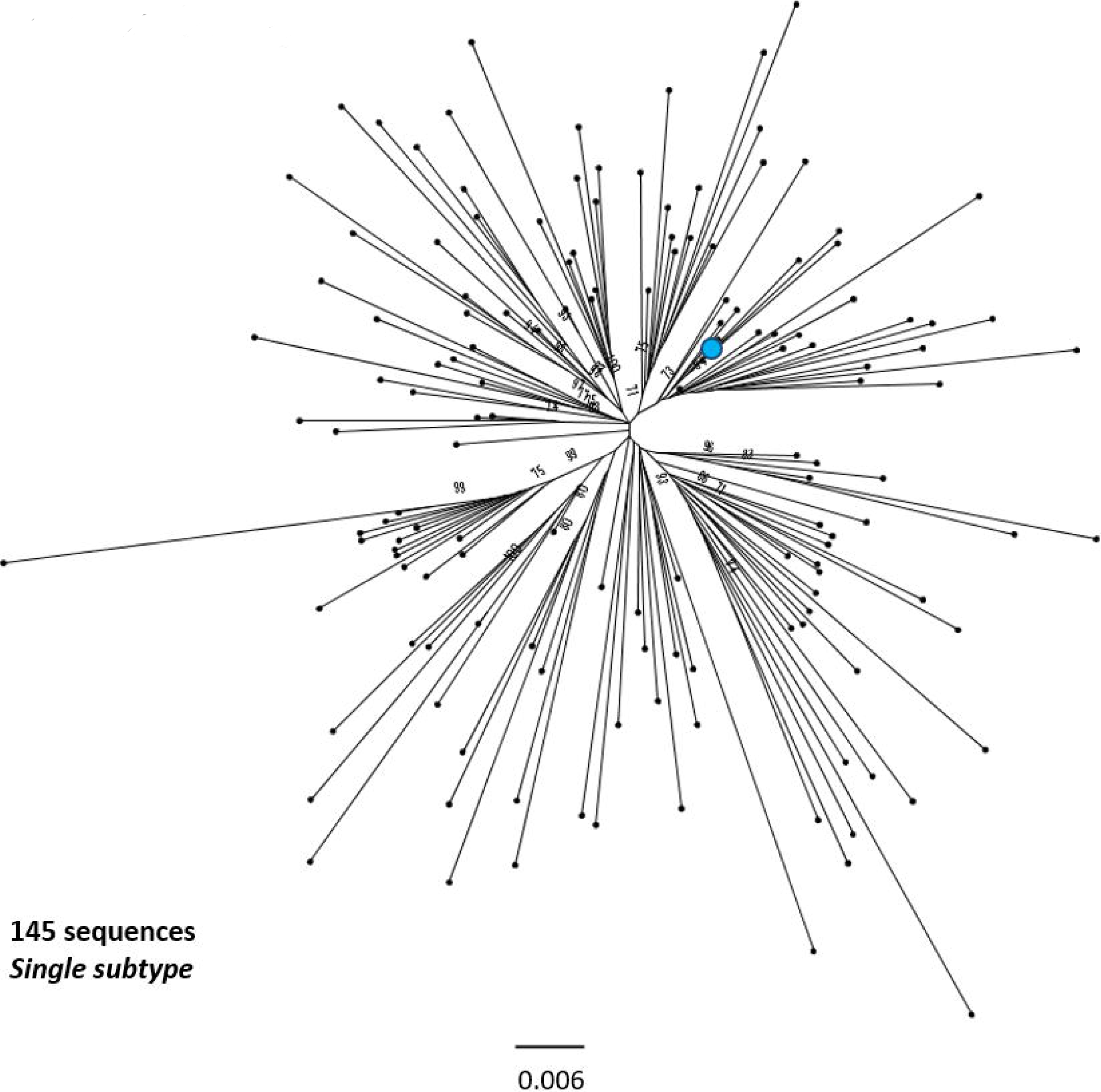
Genomic-length maximum-likelihood phylogeny of HBV genotype E sequences. Genotype E sequences do not diverge into distinct subgenotypes. Bootstrap support for branches ≥70 after 1000 replicates is indicated. The scale bar indicates the estimated nucleotide substitutions per site. Proposed reference strain for the genotype, GQ161817is highlighted with a blue dot.

**Figure 7.**
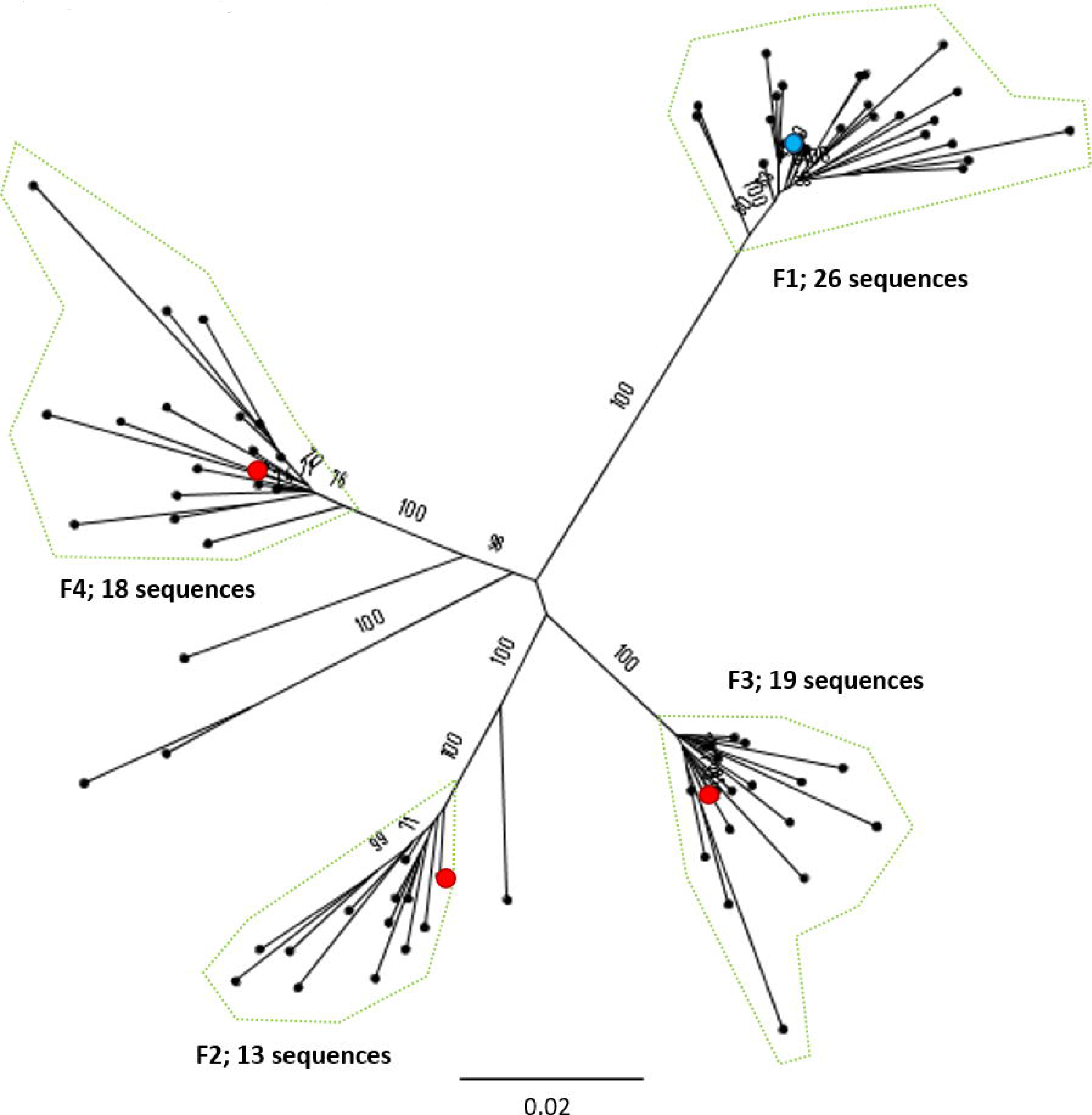
Genomic-length maximum-likelihood phylogeny of HBV genotype F sequences. Well-defined clades have been highlighted with coloured dotted lines and reference sequences for each clade indicated (red dots). Proposed reference strain for the genotype, HM585194 is highlighted with a blue dot. The subgenotype is given where it could be reliably identified. Bootstrap support ≥70 after 1000 replicates is given. The scale bar indicates the estimated nucleotide substitutions per site.

**Figure 8.**
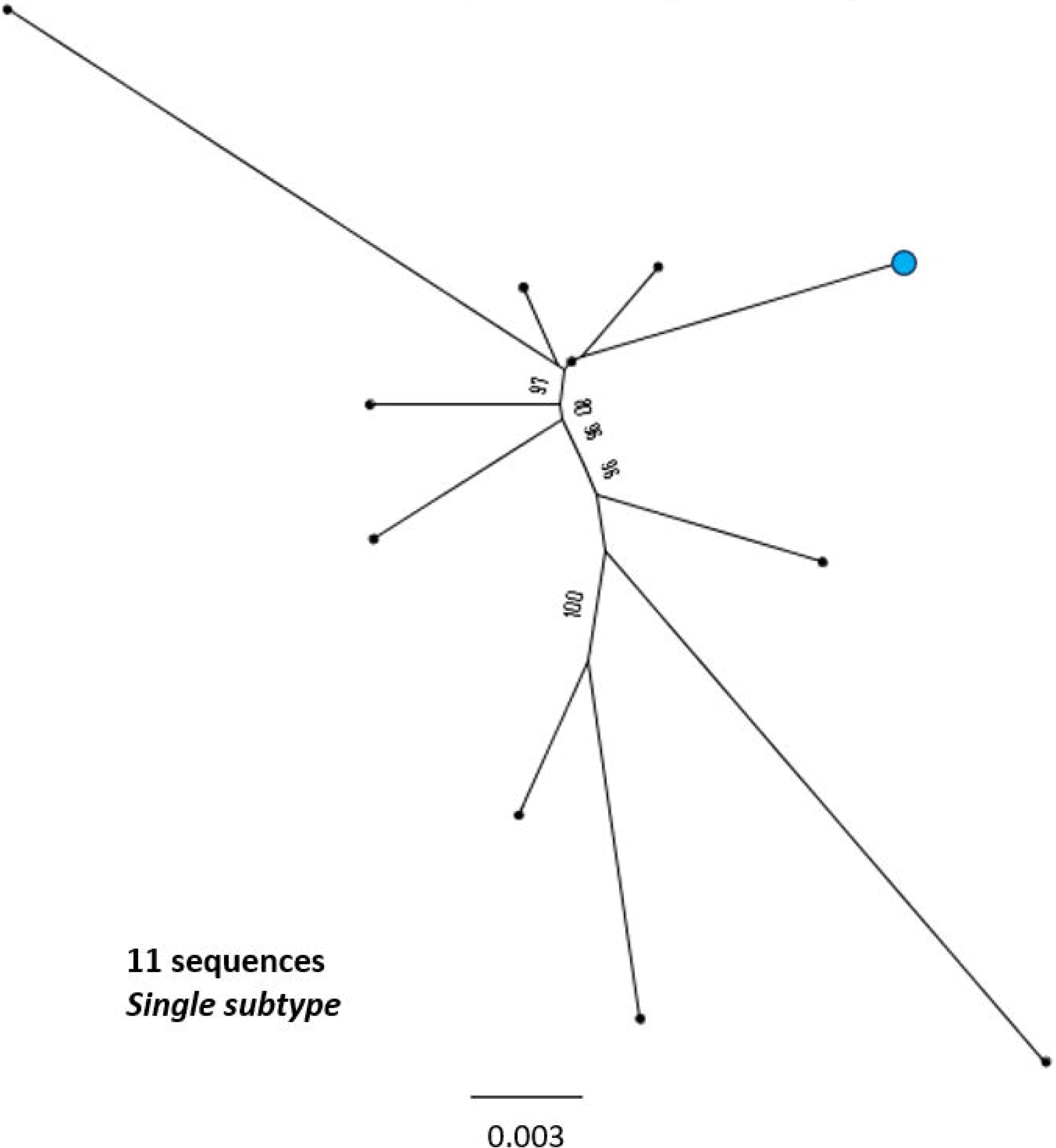
Genomic-length maximum-likelihood phylogeny of HBV genotype H sequences. Genotype H sequences do not diverge into distinct subgenotypes. Bootstrap support ≥70 after 1000 replicates is given. Proposed reference strain for the genotype, FJ356715 is highlighted with a blue dot. The scale bar indicates the estimated nucleotide substitutions per site.

**Figure 9.**
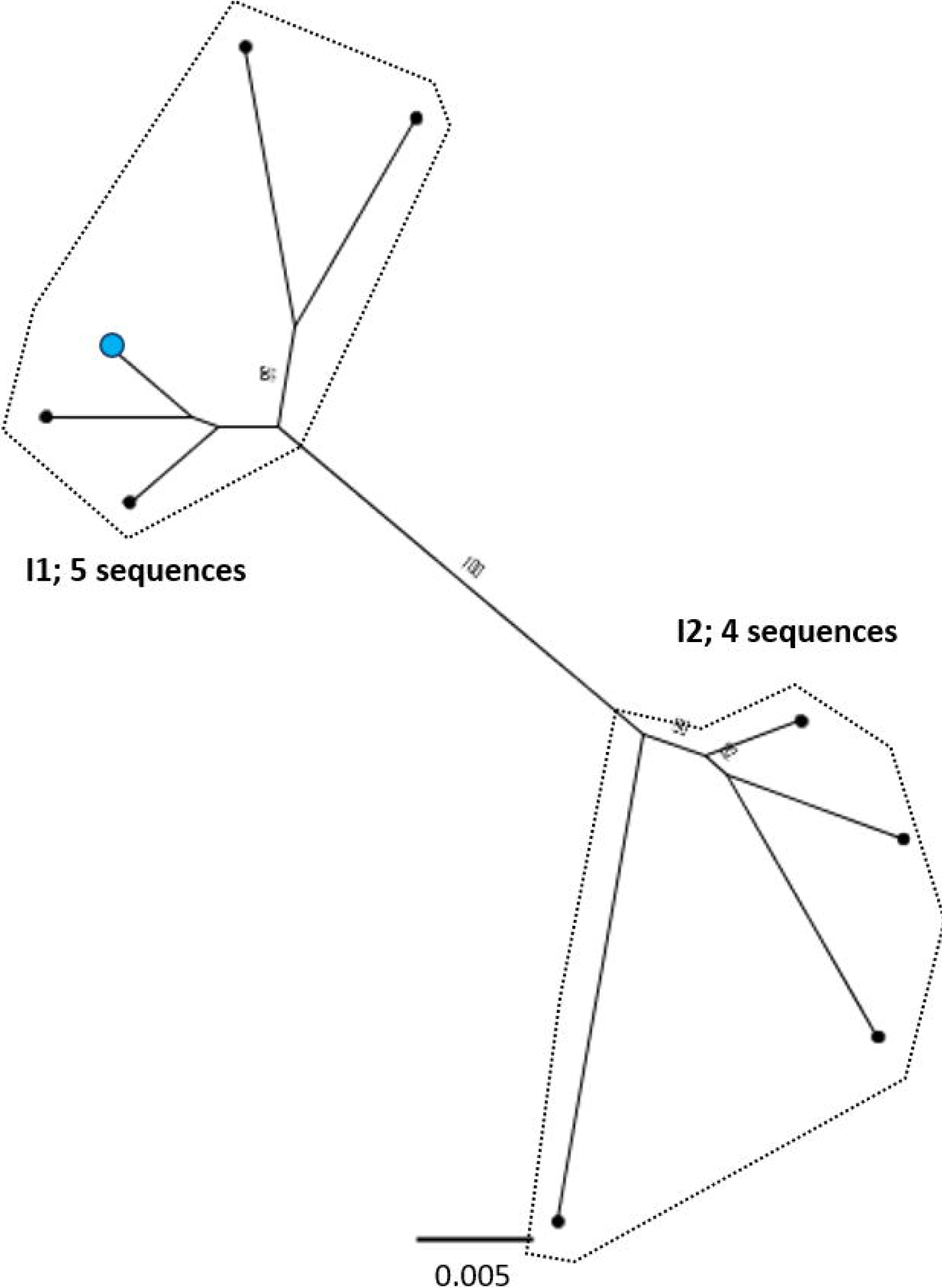
Genomic-length maximum-likelihood phylogeny of HBV genotype I sequences. Well-defined clades have been highlighted with coloured dotted lines and reference sequences for each clade indicated (red dots). Proposed reference strain for the genotype, AB562463is highlighted with a blue dot. Bootstrap support for branches ≥70 after 1000 replicates is indicated. The scale bar indicates the estimated nucleotide substitutions per site.

### Ambiguities and inconsistencies in existing data

There are a number of poorly characterised and disputed subgenotypes in the literature, particularly within genotype C, making accurate identification of the subgenotypes challenging. Strong evidence to support a number of previously described clades was also difficult to ascertain, with specific difficulties in assessment of certain subgenotypes; D6, C7 and C9 were not easily distinguishable in this analysis. A search of Genbank indicated a single genomic-length sequence of subgenotypes A4 (KM606737), A6 (GQ331046), and D6 (KF170740) in the database, and no evidence of genomic-length sequences for subgenotype C7 or C9 isolates. Both A4 and A6 isolates clustered away from other sequences in the analysis, but few additional isolates of the subgenotypes were retained after removing highly similar sequences. In the case of subgenotype C7, previous publications on the phylogeny of genotype C used EU670263 as a C7 reference strain (17,18). However, in Genbank, EU670263 is listed as subgenotype C6 (19), and clusters within our phylogenetic analysis with GU721029, which is also designated subgenotype C6. A single provisional isolate of subgenotype C9 exists (AP011108), having been proposed by Mulyanto et al. in 2010 (17) but this designation is not present in the Genbank data associated with the sequence. Likewise, subgenotypes C13-C16 have been described (20), but we were unable to distinguish these as distinct subgenotypes in our analysis. The D6 isolate KF170740 was not retained in the sequences we selected for analysis, suggesting it is closely related to another genotype D subgenotype.

### Comparative phylogenies and pairwise distance of HBV genotypes

We aligned our newly defined genotype and subgenotype reference sequences, and used these to generate a ML phylogenetic tree (Fig 10). Pairwise distance analysis for the majority of genotypes (Fig 11) revealed a bi-modal distribution of the distances, with one peak representing the relationship to sequences from the same (or closely related) clades (typically showing 1-3% divergence) and the other peak being characteristic of distance to sequences in other clades from other subgenotypes (typically 3-8% divergence). Genotype D differed from this distribution with a single peak where the majority of pairwise distances were between 2-4%. This indicates that genotype D phylogeny does not contain long evolutionary branches separating clades from each other. Genotypes E, F, H and I contained smaller number of sequences, which can impact the pairwise distances distribution. In this group genotypes E, H and I showed a uni-modal distribution of the pairwise distances. The lowest pairwise divergence was observed in genotype H, where the majority of pairwise distances were <3%. Genotype F demonstrated the greatest between-sequence diversity, where a high proportion of pairwise distances were >5%, likely a reflection of the long evolutionary distance between subgenotypes. Genotype G had only three sequences after our initial filtering which was too small for a meaningful pairwise distance analysis.

**Figure 10.**
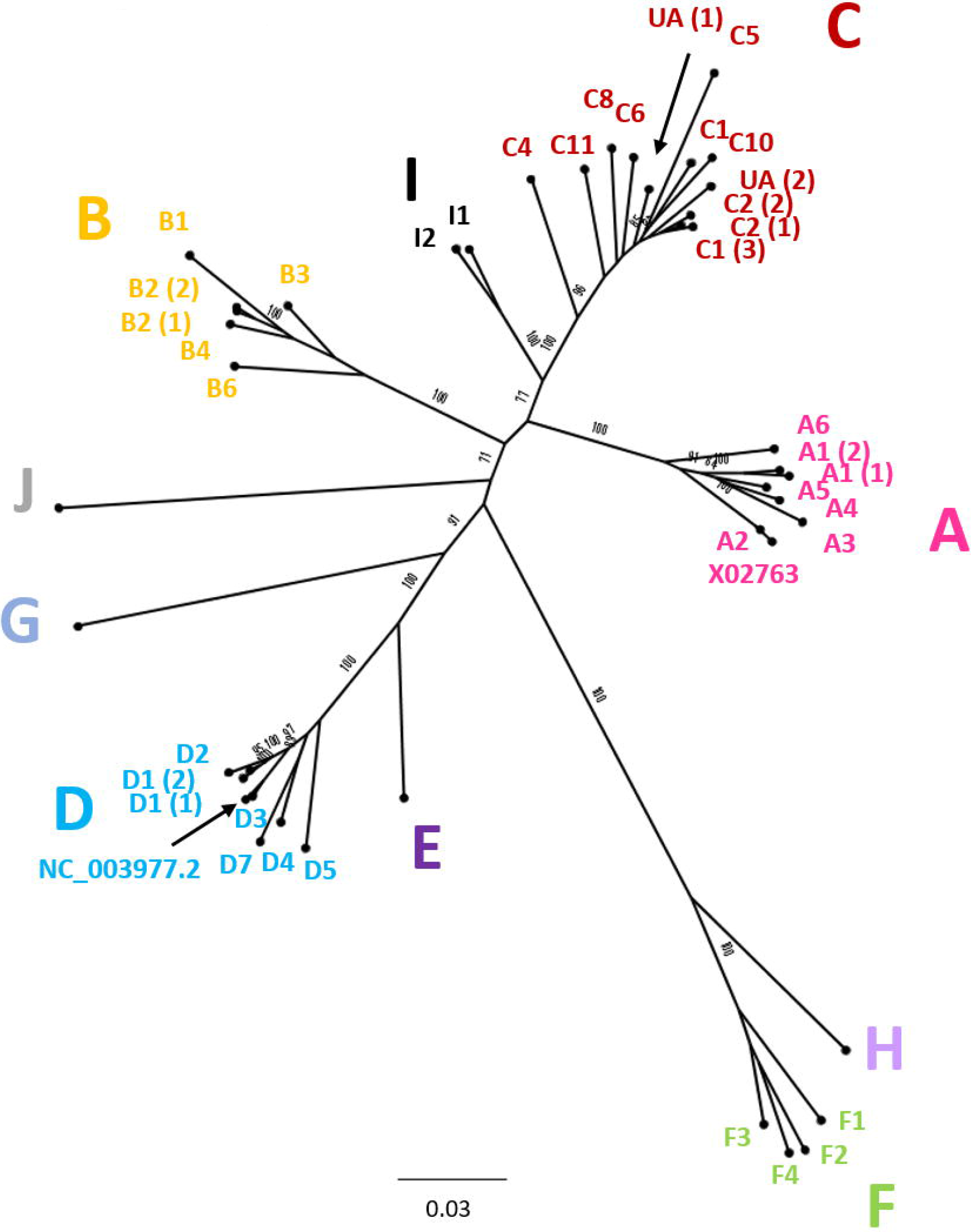
Genomic-length maximum-likelihood phylogenetic tree of HBV genotype, subgenotype and clade reference strains identified in Figures 2–9 and listed in Table 1. The genotype is given in each case and the subgenotype or clade identification is given where possible. Bootstrap support for branches ≥70 after 1000 replicates is indicated. The scale bar indicates the estimated nucleotide substitutions per site. In addition to the references identified in Figures 2–9, genotype A isolate X02763 and genotype D isolate NC_003977.2 have been included in the tree.

**Figure 11.**
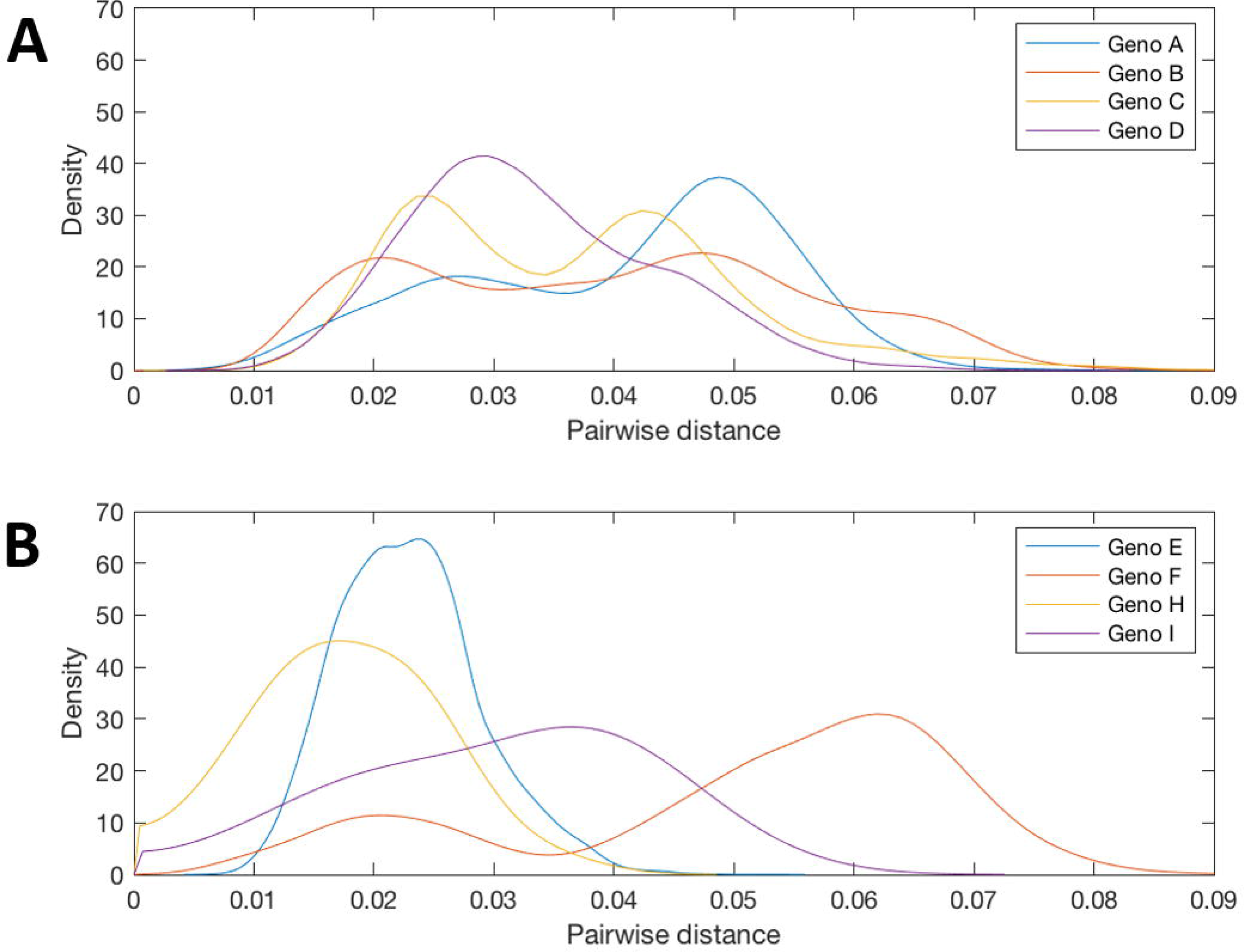
Pairwise distances distribution for the genomic-length sequences of HBV genotypes A, B, C, D, E, F, H, I. Probability densities of pairwise distances for whole genome sequences of HBV genotypes. Genotypes E, F, H and I are shown on a separate plot from genotypes A-D as they contained smaller number of sequences. Too few sequences were available after filtering for genotypes G and (putative) genotype J to be analysed.

## DISCUSSION

### Summary of this resource

We have collated genomic-length HBV reference sequences into a repository, generating a resource that will help to underpin research. As clinical practice evolves to incorporate recommendations pertaining to HBV genotype, our dataset will also potentially become useful to support sequence-based insights for clinical practice.

### Insights into diversity and geography

Genotype C has a particularly extensive phylogeny relative to the other genotypes. Genotype C is prevalent in the Asia-Pacific region and is thought to be the oldest extant genotype (21). The expansive diversity within this genotype is likely a consequence of its long evolutionary history in humans, although it should be noted that this genotype is also the most well represented genotype in the analysis, accounting for over 1/3 of all sequences in our dataset. Considerable diversity is also observed with genotypes B and D. Genotype D has a relatively broad geographic prevalence, being found in regions including the Mediterranean, North-Eastern Europe, India, Oceania and parts of southern Africa (22). It is possible that persistence of chronic HBV infection in these varied ethnic groups has contributed to the diversification of this genotype.

Genotype F has an atypical phylogeny compared to other genotypes, comprising four subgenotypes separated by long evolutionary distances, but with little intra-subgenotype diversity. It is unclear if the unusual phylogeny is a result of the elevated substitution rate documented for genotype F (23) or if there is a paucity of sampling relative to the other genotypes. The genotype is distributed throughout Latin America and the Arctic Circle (24,25), regions that are under-represented by clinical and research cohorts to date. Genotypes E, G and H, each of which has a single subgenotype, have relatively constrained phylogenies, showing low levels of genetic diversity in the currently sampled isolates. Genotype G has a particularly low evolutionary rate, likely resulting from the 36bp insertion in the core gene hampering viral replication (26). The genotype is typically observed in co-infection with either HIV or HBV genotype A2 (26).

A HBV strain isolated from a 16^th^ century skeleton (LT992440) clustered between subgenotypes in genotype A. The isolate falls (with long branch lengths) together with two sequences both isolated from patients in Belgium, one of which has been proposed as the hypothetical subgenotype A6 (GQ331046), isolated in 2006 (27). The clustering of these three sequences highlights the protracted association of HBV with human populations, and the marked lack of temporal population structure displayed by the viruses, frequently confounding phylogenetic attempts to understand the evolutionary history of HBV (28,29).

### Limitations

#### (i) Errors and inconsistencies in existing classification system

There are a number of poorly characterised and disputed HBV subgenotypes in the literature, particularly within genotype C. A cause of these inconsistencies is the assigning of ‘provisional’ subgenotypes, based on a single isolate, as is the case with C7 and C9. Using the published literature to inform our analysis, we took an indication of strong bootstrap support, and a threshold of ≥2 sequences to define and identify the subgenotype clades. For some subgenotypes, including A4 and A6, multiple isolates have been reported in the literature but few isolates were retained in our analysis after removing highly similar sequences, suggesting divergence within the subgenotype is limited. Despite this approach, a number of clades grouped distinctly away from known subgenotypes but could not be categorically assigned to a subgenotype (thus we designated two ‘unidentified clades’ in genotype C).

We suggest that, in future, a minimum number of epidemiologically unlinked sequences and genomic coverage (e.g. ≥2 genomic-length unlinked sequences) should be required to designate a new HBV subgenotype. Bias introduced into databases from intra-host diversity studies, where multiple sequences are isolated from the same patient (such as clonal analysis) can confound such analyses, but should have been largely controlled for in this case by stripping the alignments of similar sequences.

The phylogenetic methods selected may affect the subgenotypes classified, as there may be variability in bootstrap support between methods. Both neighbour-joining and parsimony methods are considered less accurate than maximum-likelihood approaches (30,31), as used in this study. Future analysis to define HBV subgenotypes should be based on maximum-likelihood methods and the bootstrap support should be indicated.

#### (ii) Bias in published data

There is considerable variation in the numbers of sequences available for each genotype, with genotypes B and C particularly well represented and genotypes E-I relatively neglected. Whilst these differences may reflect genuine variation in the contribution of different genotypes to the total global pool of infection - with B and C coming from high-prevalence, densely populated regions - significant under-sampling of HBV sequences from other regions is likely (especially from low-income regions). Similar trends in isolate bias have been reported in the HCV field (32). Consequently, the sequences available in publicly available databases are likely to under-represent the true extent of HBV diversity and, as more sequences are generated reference sets will need to be reviewed.

Several previous studies publishing HBV reference sequence sets have generated consensus sequences from alignments of full genome HBV sequences and submitted these new consensus sequences to Genbank (33,34), which leads to the potential inference that these are sequences of biological origin. A number of genotype B, D and F sequences generated with the same approach have also been submitted to Genbank but they are not supported by an associated publication, making it impossible to assess their provenance. The methods used to generate these sequences mean that, by definition, they occupy a basal position in the subgenotype clades after phylogenetic analysis and appear to be ideal reference sequences. However, these consensus sequences may differ from biologically derived sequences at key sites, and are therefore potentially misleading if used as reference sequences. Improved sequence metadata in the records of published sequences, clearly indicating the methods used to derive these sequences would be informative for researchers.

#### (iii) Recombination and ‘quasi-subgenotypes’

Recombination is relatively common amongst HBV isolates (11,35) and a number of established HBV subgenotypes have been shown to have a recombinant origin, particularly in genotypes A, C, D and the putative genotype J (35–37). A number of sequences in our phylogenetic analysis were not easily classified, as they fall phylogenetically between two well-defined subgenotype clades. These sequences are possibly inter-subgenotype recombinants, combining genetic regions from two or more subgenotypes, making them challenging to classify in genomic-length genome phylogenies.

Previous work has indicated that some designated subgenotypes, including A3, A4, A5, C2 and B3 group as distinct, monophyletic clades in maximum-likelihood analysis but that they do not meet the required genetic distance to be classified as separate subgenotypes (3,9,38). It has been suggested these should instead be designated ‘quasi-subgenotypes’ to reflect their divergent nature but underscore that they do not meet the technical definition of a subgenotype (9). In our analysis, quasi-subgenotype A3 grouped into A3, A4 and A5 clades with strong bootstrap support. It was not possible to separate subgenotypes C2 and quasi-B3 (encompasses B3 and B5) into distinct subgenotypes.

## Conclusion

Sequencing is increasingly utilised in the clinical management of HBV infection to inform on prognosis, treatment choice, drug resistance and to provide insight into complex cases (39,40). Whilst progress in applying new sequencing technologies to HBV has been slow to date, we are now entering an era of rapid change. A consistent, unified approach to classification will advance this field, improving consistency of curating, archiving and reporting sequence data. Recent work sampling poorly accessed populations has uncovered numerous new HCV isolates (41,42), and the diversity of sequences generated for HBV may expand in a similar way. The use of a robust and consistent classification system will also be important in screening new populations for potentially novel HBV isolates. All of the generated data, including the alignments, phylogenies, consensus sequences and chosen reference sequences are available online as a simple and open-access resource (https://doi.org/10.6084/m9.figshare.8851946). We have generated a new data resource for researchers and clinicians in the HBV field, providing a solid foundation for analysis, and a structure on which to build as more data emerge.

## METHODS

### Definitions

We based our analysis on previously agreed definitions regarding classification of HBV sequences on the basis of their nucleotide sequence diversity (3), as follows:

- ***Genotype:*** nucleotide divergence of >7.5% has been proposed as the threshold for the definition of distinct HBV genotypes (43).
- ***Subgenotype:*** Genotypes are further classified into subgenotypes, based on a divergence of 4-7.5%, monophyletic clustering, and strong bootstrap support for the clade (3).

In this study, we used strong bootstrap support (≥70) and monophyletic clustering to identify distinct clades and examined the pre-existing literature for the likely subgenotype designation. In instances when subgenotypes can be seen to split into distinct clades but genetic distance and poor bootstrap support do not suggest a unique subgenotype, such as in the case of subgenotypes A1 (Fig 2) and C2 (Fig 4), we have referred to these as ‘A1 (1)’ and ‘A1 (2)’.

### HBV genome numbering convention

As HBV is a circular virus, there is technically no ‘start’ or ‘end’ to the genome. We have followed convention in the field (and HBVdb) for this study, defining nucleotide (nt)1 at an *EcoR1* restriction site in the Pol/Surface overlap (GAA/<u>T</u>TC, with nt1 starting at the first T) (2). The *EcoR1* site is hypothetical in many HBV sequences, with GAA/CTC being more frequent in many genotypes.

### Sources of sequence data

We downloaded all available genomic-length genotyped HBV sequences (N=7108) for genotypes A-H from HBVdb (14) on 31st of January 2019, and removed sequences indicated as recombinant by the database (N=767), giving a total of 6341 sequences. We used HBVdb as the database runs sequences downloaded from Genbank through a genotyping algorithm prior to inclusion and all listed sequences are annotated to ensure they follow the HBV genome numbering (14), as outlined above.

We enhanced our dataset by excluding two sequences from the genotype G phylogeny, as they had been incorrectly genotyped with the HBVdb algorithm. Both sequences (FJ023674 and FR714503) were found to be partial genotype C recombinants. We also identified an additional 71 sequences belonging to genotype I and to genotype C and D subgenotypes (under-represented in the HBVdb database) on Genbank and added these to our data set, giving a total of 6412 sequences. Ancient HBV sequences, isolated from bodies hundreds of years old (28,44), are present in these databases and they were not excluded from this analysis, unless they grouped distantly from a known genotype.

### Existing reference sequences

We included the genotype A strain X02763 (45) in the alignment with other reference sequences as this sequence (length 3221bp) is widely used as a numbering reference. Only Geno-G has insertions relative to genotype A. We suggest that the current NCBI HBV Reference Sequence, NC_003977.2 (46), a genotype D isolate, is less suitable as a numbering reference as genotype D strains have a 33-bp deletion in pre-S1, making them the shortest length HBV genotype at 3182bp. Nonetheless, we also included NC_003977.2 in the reference alignment as it is frequently used as a reference.

### Phylogenetic analysis

We aligned all sequences using MAFFT (47) and calculated pairwise distances between all isolates. In order to reduce the possibility that multiple sequences from a single individual or closely related cluster could dominate the phylogeny, we kept only one sequence from each group of identical sequences, and used hierarchical clustering to group isolates based on their pairwise distance and identified clusters of sequences that were within ≤1% of each other, retaining a single sequence from each cluster for the analysis (retaining the sequence with the lowest proportion of ambiguous sites). After stripping out these closely related sequences, we generated a maximum-likelihood (ML) phylogenetic tree of the remaining 2839 sequences using RAxML (48), including 1000 bootstrap replicates (Fig. 1). We used this tree to manually delineate sequences belonging to each genotype and excluded any remaining recombinant sequences that had not been identified earlier in the process.

We aligned the sequences for each genotype separately, and generated ML phylogenetic trees as previously described. We examined ML phylogenetic trees and generated bootstrap values to define distinct clades containing ≥2 sequences with strong phylogenetic support (Figures 2–9). Phylogenetic trees were examined in both traditional and radial layouts to ensure that the subgenotypes and clades were being delineated accurately. We excluded Genotype G from the phylogenetic analysis, as after removing highly similar sequences there were just three sequences remaining.

### Definition of new reference sequences

For each well-defined clade, we generated a consensus sequence and selected the closest clinical isolate to the consensus sequence as the reference strain for that clade. We applied a process of quality assurance to selection of reference strains as follows:

- We verified that isolates selected as reference strains were associated with an associated publication indexed in PubMed;
- We checked that reference strains were genuine clinical isolates (rather than *in silico* reconstructions such as consensus sequences);
- We rejected isolates that had insertions or deletions relative to the majority of sequences in the clade;
- We avoided (where possible) sequences with many ambiguous sites;
- We avoided use of sequences containing known drug resistance mutations as reference strains. Thus in the situation in which a sequence containing an rtM204 substitution in the ‘YMDD’ motif (associated with escape from lamivudine therapy(49)) was initially selected as a reference sequence, we rejected this and selected the next closest sequence to consensus.
- Three subgenotype references were selected with Hamming distances >50 relative to the consensus sequence for the subgenotype clade (subgenotypes C4, C11 and C_UA2). Alternative, more closely related sequences were often present within the subgenotype clade but indels in the sequences and a lack of supportive publications meant that other sequences were selected.

To define genotype reference sequences, we selected the reference sequence for the subgenotype that contributed the largest number of sequences to that genotype (grey boxes, Table 1). In cases where there were multiple clades for the subgenotype, we selected the sequence from the largest clade as the reference. Where there was strong evidence to support specific subgenotypes from both the literature and the phylogenetic analysis, we included the subgenotype in the analysis. Where subgenotype designation was unclear, we used BLAST to compare the sequence(s) to other sequences in online databases to see if the subgenotype could be identified. We also examined trees to find the accession numbers of subgenotypes that did not have an easily identifiable clade in the analysis using isolates listed in Genbank. If they were not present, these subgenotypes were most likely removed during the pairwise analysis for being highly similar to sequences already in the analysis, suggesting further examination of their designation as a unique subgenotype may be warranted. Where the subgenotype could not be identified, the clade was labelled ‘unassigned’.

In situations in which we could not ascertain which subgenotype a clade belongs to, despite examination of Genbank records and BLAST analysis of sequences, we have described these as ‘unassigned clades’ (both in genotype C, Fig 4).

## Declarations

### Authors’ contributions

AM and PM conceived the idea for this project. AM and MA identified and analysed the sequences in this study. AM, MA and PM wrote the manuscript and all authors read and approved the final manuscript.

### Availability of data and material

The data described in this Data Note, including the genotype alignments (fasta format), consensus sequences and, phylogenetic trees (newick format) are available in an open access format on Figshare (https://doi.org/10.6084/m9.figshare.8851946).

### Ethics approval and consent to participate

Not applicable

## Acknowledgements

Not applicable

