## Supplementary file for "Analysis of genomic-length HBV sequences to determine genotype and subgenotype reference sequences"

**Supplementary figure 1**; **Cumulative number of whole genome and partial HBV sequences in Genbank, 2000-2017.** Data downloaded directly from Genbank in January 2018.

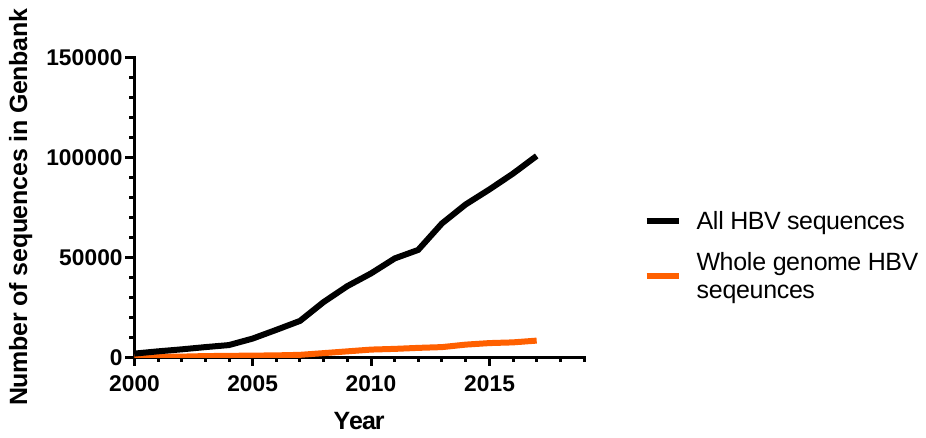
